# OpenStride: an inexpensive, open-source force plate actometry system for quantification of rodent motor activity and behaviour

**DOI:** 10.64898/2025.12.17.695041

**Authors:** Yang Yang, Brock Cooper, Michael Houghton, Kajal Dahiya, Sarah E Haslam, Henry Blackwell, Kaanchana Sekaran, Lingye Hu, Omid Kavehei, Collin J Anderson

## Abstract

**Introduction:** Force plate actometry (FPA) enables quantitative assessment of rodent motor and behavioural activity. Through tracking centre of mass at high temporal and spatial resolution, FPA can directly quantifying motor performance and various behaviours in the context of operator-independent, naturalistic movement. Previously designed systems have been expensive and non-customisable, and they are no longer commercially available.

**Methods:** We designed OpenStride, an open-source FPA system consisting of both hardware and software. Our system was designed to be able to be constructed with standard tools worldwide at a cost of ∼$550 USD, with modifiable hardware files, modular software, and support for both Mac and Windows operating systems. Required tools include 3D printing and acrylic laser cutting.

**Results:** OpenStride reliably tracks position within a 30 × 30 cm^2^ environment. Based on these positional data, OpenStride currently quantifies tremor, ataxia, distance, velocity, low-mobility bouts, and centre-*vs*-margin time, with potential to expand to additional analyses.

**Conclusions:** OpenStride is intended to provide a valuable tool for high-throughput, inexpensive study of motor and behavioural function and dysfunction. Software and hardware files will be freely disseminated online via GitHub at the time of final publication to enable others to construct and utilise the OpenStride system.

## 1 Introduction

Force plate actometry, first published on in 2001, generates the ability to track and analyse naturalistic movement and behaviour in rodents with high spatial and temporal precision (Fowler et al., 2001). A typical force plate actometer (FPA) consists of a stiff platform mounted on four force transducers or load cells; as the subjects ambulates through the chamber, its centre of mass is calculated by the weighted averages of forces applied to each transducer. Coupled with both real-time and post-experimental analyses, this enables the study of a variety of simple and complex behaviours and movements across a large variety of applications (Zarcone, 2022).

Since its initial publication, force plate actometry has been used to study numerous models of various disorders, particularly in the context of neuroscience, psychology, pharmacology, and biomechanics (Figueroa et al., 2023; Godar et al., 2016; Hein et al., 2012; Kuo et al., 2019; Lang et al., 2020; Levant et al., 2011; McCarson et al., 2019; Mosher et al., 2019; Mosher et al., 2018; Scoles et al., 2022; Zarcone et al., 2009), spanning several species, including mice, rats, voles, pigeons, and humans (Crosland et al., 2005; Pinkston et al., 2008; Tickerhoof et al., 2020; Zarcone, 2022). Across this range of models and species, specific measurements made have included distance travelled, rearing, neurodevelopmental phenotypes, operant conditioning, exploration of novelty, stereotypies, ataxia, and tremor (Fowler et al., 2003; Fowler et al., 2010; Levant et al., 2011; Levant et al., 2010; Zarcone, 2022).

We have utilised force plate actometry repeatedly in recent years. First, we combined hardware modified from that initially developed by Fowler with custom software to quantify ataxia, tremor, and falls in the Wistar Furth *shaker* rat, before evaluating deep brain stimulation targeting the deep cerebellar nuclei for degenerative cerebellar ataxia and tremor (Anderson et al., 2019; Lang et al., 2020). Next, we utilised a similar setup to perform automated open-field experiments, studying the effects of *Stau1* knockdown in the context of targeting this gene therapeutically to treat amyotrophic lateral sclerosis and spinocerebellar ataxia type 2 (Scoles et al., 2022). Subsequently, we used force plate actometry to demonstrate changes associated with functional complementation to prove causality of *Slc9a6* mutation in the *shaker* rat, linking it to Christianson syndrome (Figueroa et al., 2023). More recently, we have combined actometry with machine learning to automate the detection of complex grooming behaviours (Anderson et al., 2024). Finally, we utilised actometry to evaluate viral gene therapies for Christianson syndrome in the *shaker* rat (Anderson et al., 2024). Collectively, this body of work has led us to find actometry to be a highly versatile and reliable approach for sophisticated behavioural and motor quantification.

Previous generations of force plate actometers have several limitations. First, systems are no longer produced or commercially available. Second, costs remained relatively high, limiting uptake in a high-throughput manner. Third, given proprietary software, open-source capabilities to add analyses were not readily present. Thus, we sought to design a novel, inexpensive, open-source force plate actometry system. In this work, we report on our efforts to develop a novel force plate actometry design based on the goals of minimising cost, designing software and hardware in an open-source fashion, ensuring easily reproducible manufacturing, and maximising reliability. We also report on experiments designed to test precision and reproducibility. To this end, we sought to utilise readily available or easily manufactured components, primarily those that can be assembled on inexpensive 3D printers, while also creating open-source software that can be used with both Mac and Windows operating systems. In total, our cost to produce the new force actometer was approximately $550 USD. Thus, at the point of final publication, we will freely post all documentation, schematics, and software needed to replicate our design with tools available at a standard university fabrication laboratory.

## 2 Methods

### 2.1 Animals

All animal work was completed under two University of Sydney Animal Ethics Protocols. In total, one adult mouse and 13 adult rats were utilised in this work. In both instances, rodents were housed 2-5 per cage under a standard 12:12 hour light cycle and given access to food and water *ad libitum*. The mouse utilised was of the C57BL/6 background, and the 13 rats utilised were Wistar Furth, including 7 wild-type and 6 *shaker* rats hemizygous (male) or homozygous (female) for a loss-of-function mutation in the *Slc9a6* gene, which yields progressive cerebellar degeneration and ataxia (Figueroa et al., 2023).

### 2.2 Hardware and Software Architecture

The OpenStride system consisted of a rigid 3D-printed base supporting a 30 × 30 cm^2^ 3D-printed walking surface enclosed within a 5-mm acrylic chamber. Four single-point load cells (Tedea Huntleigh Model 1004-00.3-JW00-RS) were mounted beneath the walking platform in a square configuration. STL files for all printed components, acrylic cutting templates, and assembly schematics will be released GitHub at the time of final publication. A 35 × 35 × 5 cm concrete paver was positioned beneath the base, on top of ethylene-vinyl acetate (EVA) foam, under each corner to shield the system from external perturbation and improve its mechanical stability.

Each load cell transmitted analogue differential signals to a PhidgetBridge 1046_0B interface, which relayed digitised voltage values to a computer via USB. Real-time acquisition was implemented in Python using the Phidget21 API, with a maximum sampling frequency of 125 Hz. Raw voltage values from each channel were buffered in system memory and written to disk in 20-s intervals using an SQLite-based routine. Timestamps were generated using the local system clock, and sequential buffer indexing was used to maintain consistent ordering of samples across write cycles.

A MATLAB App Designer interface was developed for experiment control, including configuration of sampling rate, output directory, and metadata fields. Prior to each recording session, the interface triggers a Python-based zero-load calibration script to align baseline offsets across the four load cells. Real-time visualization of the centre-of-mass (COM) position was provided by the Python acquisition program once recording commenced. Raw voltage signals were stored without acquisition-stage filtering to support analyses requiring unprocessed force traces, such as tremor quantification. A complete list of hardware and software resources required to reproduce the system is provided in Table 1, and a hardware schematic is shown in Fig. 1.

**Table 1.**
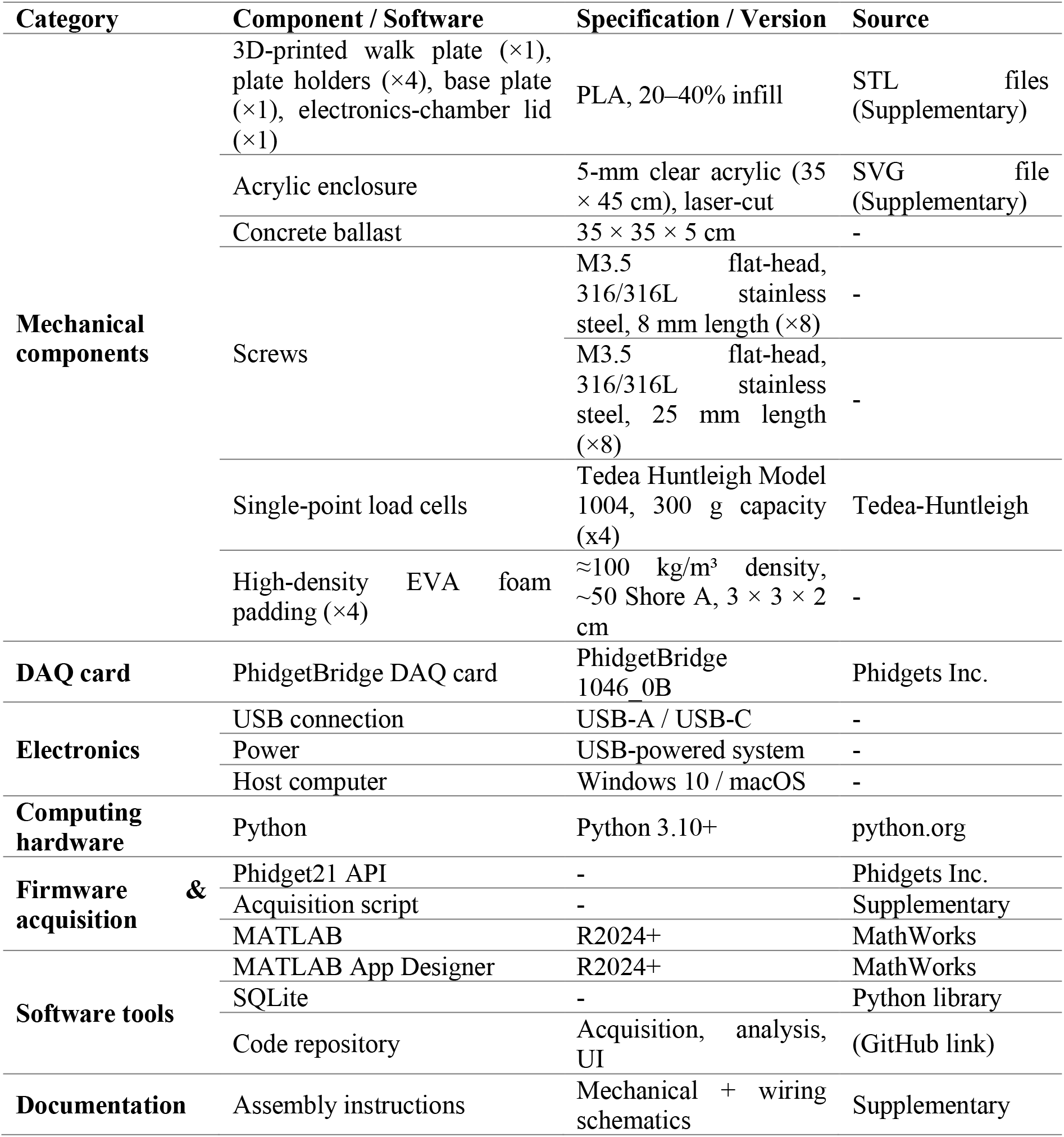
Key resources for the production of the OpenStride system

**Figure 1:**
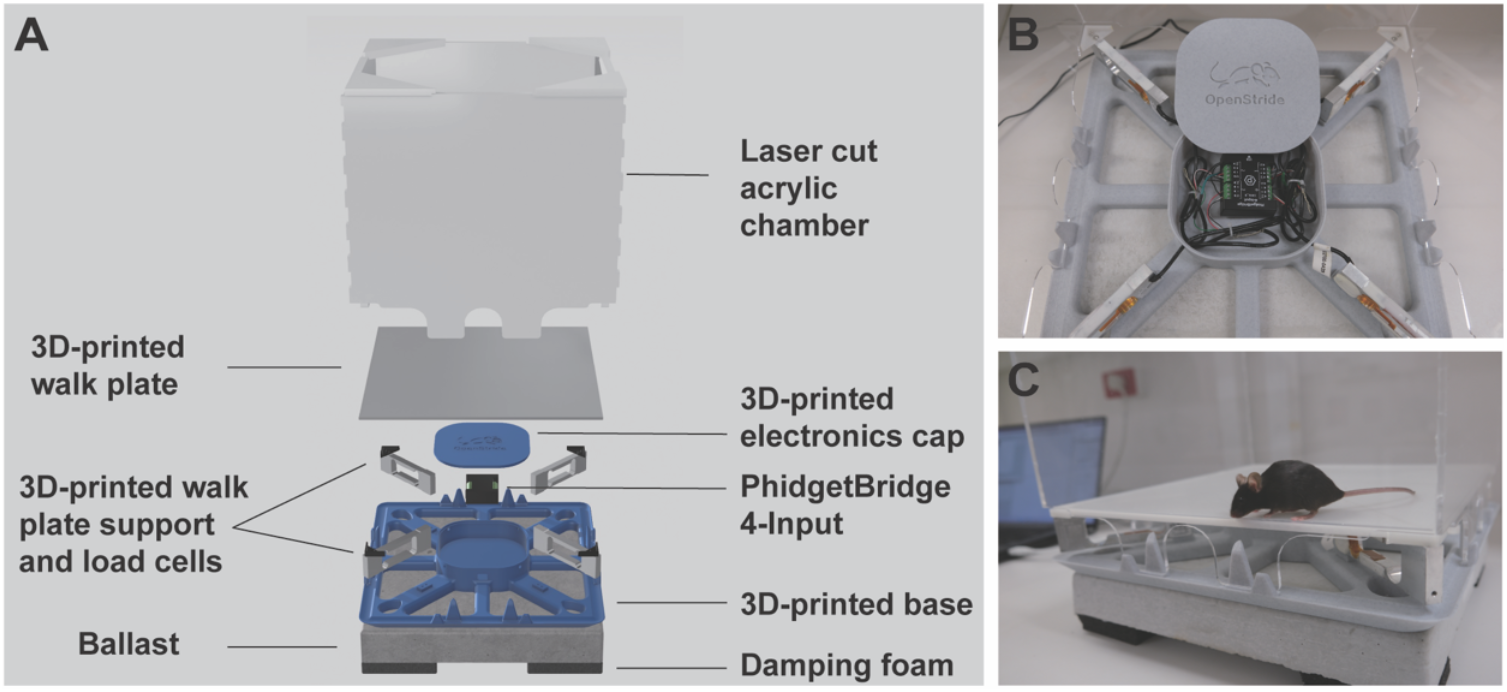
**A**. Primary components of OpenStride hardware. **B**. Assembled OpenStride without the walking plate and with the electronics lid removed. **C**. Fully assembled OpenStride system with a C57BL/6 mouse for scale.

### 2.3 Mechanical stability validation

The mechanical stability of the OpenStride system was evaluated using a series of bench-top tests performed without animals, based on computed COM trajectories obtained during acquisition. For all tests, force signals were sampled at 125 Hz.

To characterise static drift, a 100 g calibration mass was placed at the centre of the walking surface, and recordings were acquired for 60 s at 125 Hz. COM time-series data *x*(*t*) and *y*(*t*) were segmented into fixed-length, non-overlapping windows (1 s; 125 samples per window). For each window, drift amplitude was calculated as the radial standard deviation of COM position, defined as follows: *D*=var(*x*)+var(*y*). For each trial, window-wise drift amplitudes were summarised to obtain a single drift metric per recording.

To assess the effect of load magnitude on stability, the static drift protocol was repeated with calibration masses of 0, 20, 50, 100, 200, 500, and 1000 g placed at the walking platform centre. For each load condition, three independent 30-min recordings were obtained. COM trajectories were processed as above, and drift amplitude *D* was computed for each window and summarised for each trial, providing a load–drift relationship across the tested mass range.

Susceptibility to external mechanical disturbance was assessed by dropping a 2.7 g ping-pong ball from a fixed height of 10 cm onto the benchtop at 10 cm from the force-plate base. Each trial consisted of a 10-s unloaded baseline recording followed by a ball impact occurring 5-s into the recording. The disturbance window was defined as the 2-s interval following the first ball–surface contact. Peak COM displacement during this window, relative to the mean COM position during the 2-s baseline period, was used to quantify the transient effect of the disturbance on system stability.

The disturbance protocol described above was repeated with additional static loads placed at the centre of the walking surface, with masses of 20, 50, 100, 200, 500, and 1000 g tested. For each load condition, five ball-drop trials were collected. For each trial, baseline COM position was taken from the 2-s period preceding impact, and transient disturbance response was quantified as the peak COM displacement within the 2-s post-impact interval. This procedure enabled assessment of how the load on the platform influenced the magnitude of transient disturbance effects.

### 2.4 Positional accuracy validation

Distance measurement was assessed using a finger-tracing procedure performed on the walking surface. The effective working area of the walking platform was 30 × 30 cm^2^. During each trial, a finger was placed at one corner of the platform and moved diagonally across to the opposite corner before returning to the starting point; a complete forward–return motion was defined as one trace. A total of 6 traces were collected. For each trace, the measured path length was computed as the cumulative distance across consecutive COM samples: 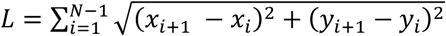. The theoretical straight-line distance between opposite corners of the 30 × 30 cm^2^ workspace was calculated as 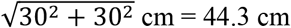.

To evaluate the effect of temporal smoothing on position, and therefore, distance, moving-average filters of varying window lengths (1–25 samples) were applied to the COM trajectories prior to distance estimation. For each smoothing condition, we quantified using the ratio between measured and theoretical distances. This procedure characterised how smoothing and sampling resolution influenced trajectory-based distance estimates across repeated diagonal traces.

### 2.5 Locomotor behaviour based on trajectories

Spontaneous locomotor activity was recorded by placing each animal individually into the hardware chamber for a 10-min session. Animals were allowed to freely explore the 30 × 30 cm^2^ arena without external stimuli. Force signals were sampled at 125 Hz and converted to real-time COM coordinates during acquisition. For analysis, COM trajectories (*x*(*t*), *y*(*t*)) were smoothed using a 15-sample moving-average filter.

Several COM-based metrics were extracted from the 10-min recordings. Path length was calculated as the cumulative Euclidean distance between successive COM samples: 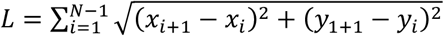. This yielded a time-resolved cumulative distance curve over the full session. Instantaneous speed was defined as the displacement between consecutive COM samples divided by the sampling interval: 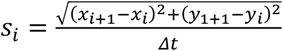.

Low-mobility bouts were identified when average speed fell below 0.5 cm/s for one second or longer. Cumulative low-mobility bout duration was computed as the sum of all time points meeting this criterion. These metrics enabled characterisation of locomotor trajectories, speed profiles, and low-mobility computations throughout the 10-min exploration period.

Times spent in the centre *vs* the margin were computed based on a defined central area. In this manuscript, the centre was defined as a square occupying the central 50% of the arena, whilst the margin was defined as the remaining 50%. This threshold is modifiable in OpenStride, and notably, may require adjustment based on subject size, i.e. rats *vs* mice.

### 2.6 Ataxia and tremor quantification

Ataxia and tremor metrics were derived from COM trajectories and speed signals obtained during spontaneous locomotion sessions. Locomotor ataxia was quantified using an ataxia ratio computed over 1-s windows. COM trajectories (*x*(*t*), *y*(*t*)) were segmented into non-overlapping 1-s windows. For each window *w*, when net displacement exceeded a pre-set threshold (10 mm used herein), cumulative path length was computed as 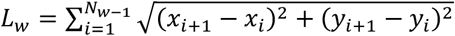, where *N*_*w*_ is the number of samples in the window. Net displacement was defined as the Euclidean distance between the first and last COM coordinates of the window 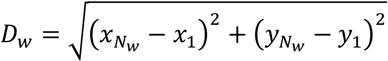. The ataxia ratio for each window was then calculated as 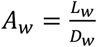. This metric thus captures the relative irregularity in trajectory, with higher ratios reflecting greater deviation from straight-line motion. Ataxia ratios were computed for all windows across each animal’s recording.

Tremor quantification was based on speed fluctuations derived from COM trajectories. Speed was obtained as *s*_*i*_ = ((*x*_*i*+1_ − *x*_*i*_)^2^ + (*y*_*i*+1_ − *y*_*i*_)^2^)/*Δt*, where Δt=8 ms is the sampling interval when sampling at 125 Hz. The speed time series was detrended and segmented into 2-s windows with 50% overlap. Power spectral density (PSD) estimates were computed using Welch’s method with a Hanning window. Tremor amplitude was defined as the integrated spectral power within a specified band (3–8 Hz used here): 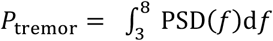. To normalise tremor magnitude across animals and trials, the tremor score was calculated as the ratio of the tremor band to the total power in the 0–20 Hz band: 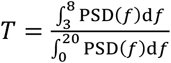. This yielded a dimensionless tremor index for each recording.

Ataxia ratios and tremor scores were computed from recordings obtained from wild-type and *shaker* rats under identical conditions. For each animal, metrics were averaged across all valid 10-min locomotor sessions. These values were used for statistical comparisons between genotypes in the results section.

### 2.7 Statistical analyses

Statistical analyses reported were performed within the MATLAB (MathWorks) environment. Comparisons between wild-type and *shaker* rats for ataxia and tremor metrics were conducted using one-tailed heteroscedastic two-sample (Welch) t-tests. Data are presented as mean ± standard deviation (SD) or mean ± standard error of the mean (SEM), with statistical significance defined as *p* < 0.05.

## 3 Results

### Mechanical stability of OpenStride

We first evaluated the mechanical stability of the OpenStride system under static and perturbed conditions. We placed a 100 g mass at approximately the centre of the walking platform for 60 s and showed that the centre-of-mass (COM) positions remained tightly clustered, with drift amplitudes below 200 μm throughout the recording (Fig. 2A). Next, we quantified how stability varied with different applied loads. Increasing the mass from 0 to 1000 g progressively reduced the drift amplitude, resulting in an approximately monotonic decrease in positional fluctuation across the tested range (Fig. 2B).

**Figure 2:**
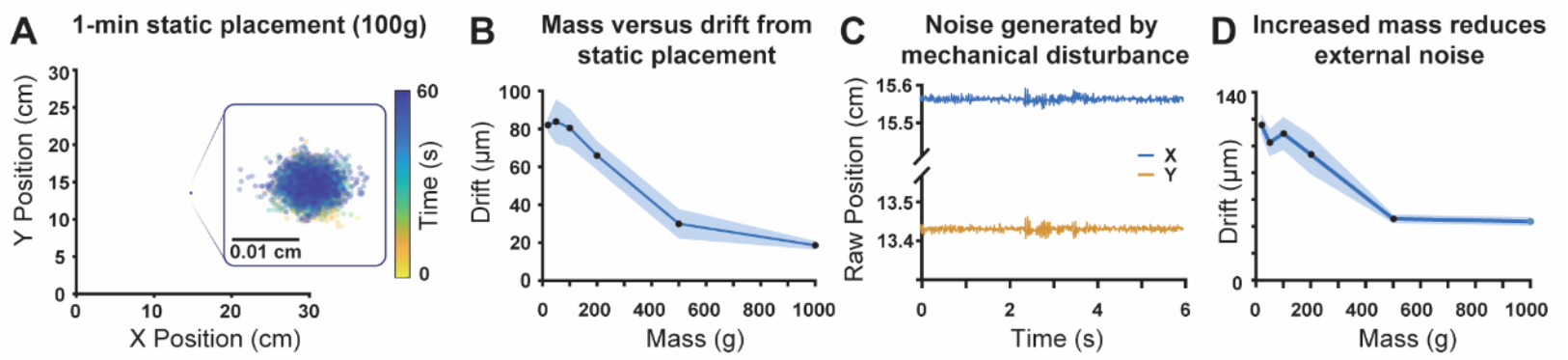
**A**. Centre-of-mass (COM) samples recorded over 60 s remain tightly clustered near the placement point. The spatial spread of COM positions indicates minimal drift during static loading. **B**. Drift decreases as load increases from 0 to 1000 g. Shaded regions represent SD. **C**. A 2.7 g ping-pong ball dropped from a height of 10 cm at a distance 10 cm from the system at ∼2 s into the recording induces a brief drift in both x and y signals. **D**. Peak transient drift decreases with load in the context of external mechanical noise.

We then examined susceptibility to external mechanical disturbance. A 2.7 g ping-pong ball was repeatedly dropped onto the benchtop from a fixed height of 10 cm and at 10 cm from the device, producing a clear transient displacement of the COM signal in the unloaded condition (Fig. 2C). Placing additional load on the platform markedly attenuated this effect: transient deviations decreased sharply with increasing load, suggesting increased stability with higher-mass subjects (Fig. 2D).

### Accuracy of position and distance

Positional accuracy was evaluated using trajectories recorded during diagonal finger-tracing movements. Each trace consisted of a forward movement from one corner of the 30 × 30 cm^2^ workspace to the opposite corner, followed by a return to the starting point. Across 20 diagonals, the recorded COM trajectories formed diagonal paths that extended across the length of the workspace (Fig. 3A). The trajectories from different repetitions followed similar spatial paths, with variations in both the x- and y-coordinates observable along the diagonals. For each trace, applying a convolutional window with a width of 10 samples, the cumulative measured distance increased approximately in step with the theoretical true distance (Fig. 3B). We then varied the size of the convolutional window. The ratio of measured to theoretical distance decreased as the window size increased from 1 to 10 samples, after which the ratio showed minimal additional change with larger window sizes (Fig. 3C).

**Figure 3:**
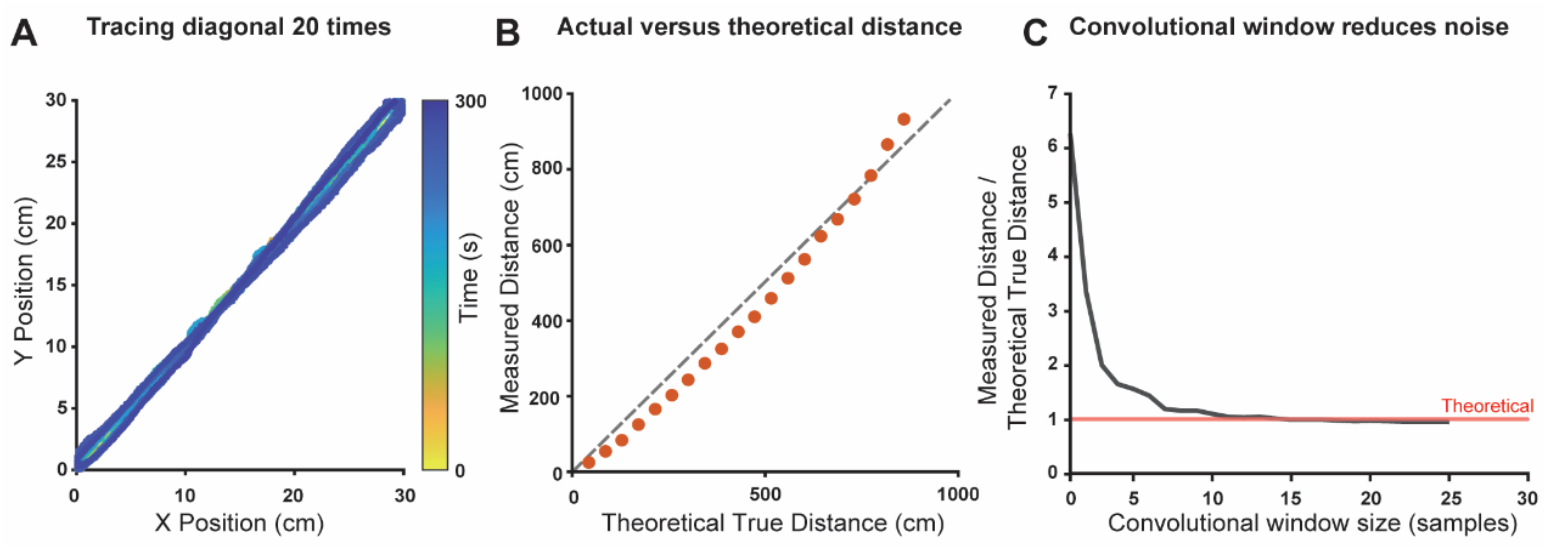
**A**. Trajectories recorded during 20 diagonal traces across the 30 × 30 cm^2^ workspace. Colours represent time within each trace. **B**. Cumulative measured distance (with a convolutional window of 10) plotted against the theoretical diagonal distance (dashed unity line). **C**. Ratio of measured to theoretical distance as a function of convolutional window size (1–25 samples).

### Wild-type rodent locomotor validation

We evaluated locomotor behaviour using COM trajectories obtained during spontaneous exploration in both a wild-type C57BL/6 mouse and a wild-type Wistar Furth rat. Representative 5-min trajectories in the 30 × 30 cm^2^ arena are shown (Figs. 4A, 4D). The wildtype mice spent most of its time in the margin, while the wildtype rat spent similar time in the defined centre *vs* margin (Figs. 4B, 4E).

**Figure 4:**
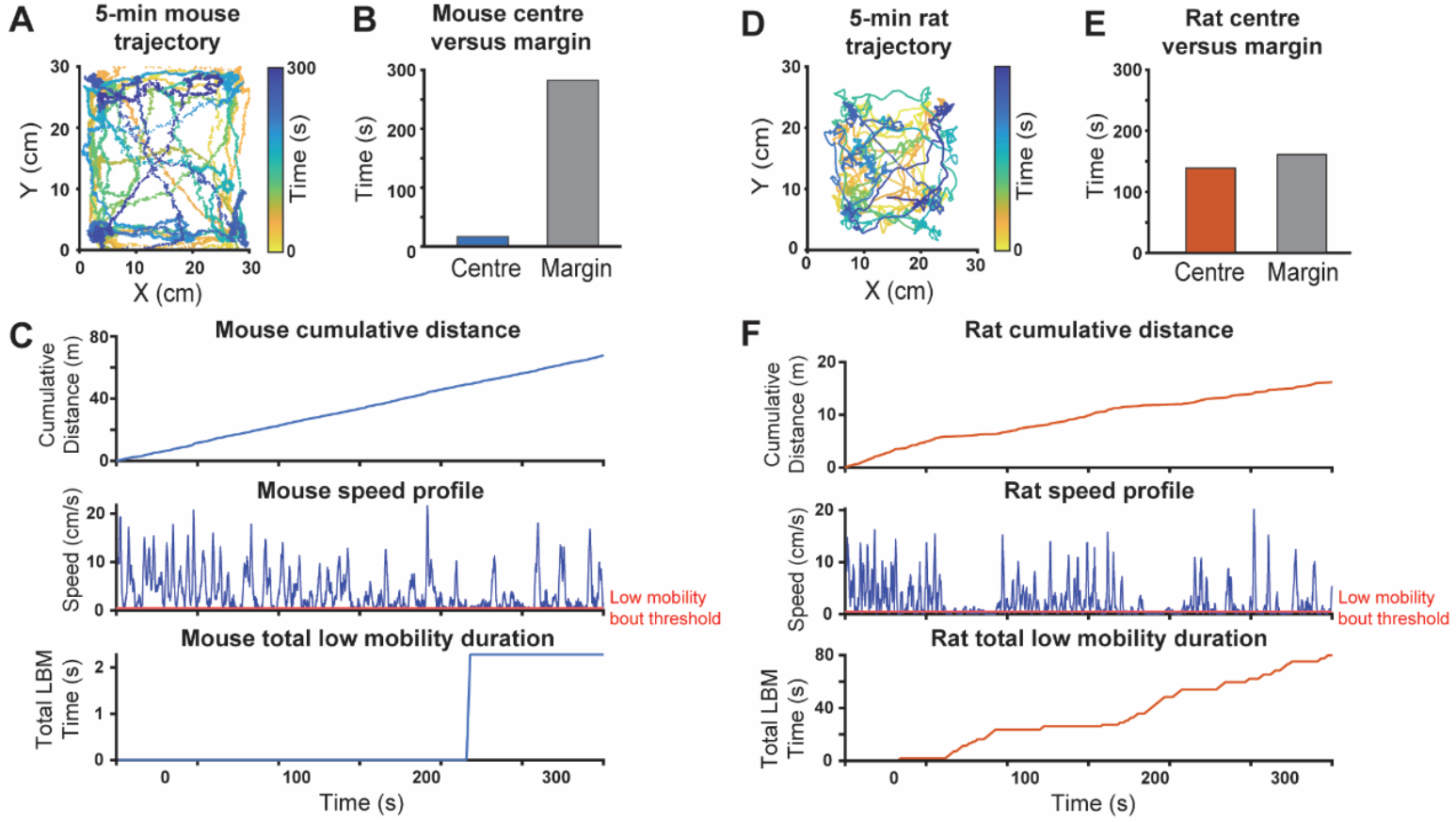
**A**. Trajectory from a 5-min mouse recording within the 30 × 30 cm^2^ workspace. **B**. The mouse spent most of its time in the margin rather than in the centre 50% of the arena. **C**. Cumulative distance, speed over time, and cumulative low mobility duration are shown from the same recording, with only one short period of low-mobility bout recorded. **D-F**. A 5-min rat recording is shown as in A-C. Notably, the rat travelled considerably less distance and had substantially more low mobility duration.

Cumulative distance, speed profile, and cumulative low mobility duration (with a 0.5 cm/s threshold applied) are shown for each 5-min session (Fig. 4C, 4E). Notably, the mouse ambulated more consistently than the rat during the 5-min recording duration, with only a single ∼2 s low-mobility bout, resulting in substantially more distance travelled than the rat.

### Tremor and ataxia measurements

Finally, we examined COM-derived ataxia and tremor metrics in wild-type Wistar Furth (n=6) and Wistar Furth *shaker* (n=6) rats. Representative 1-s trajectory segments showed a broad range of ataxia ratios, with higher ratios corresponding to increasingly irregular paths (Fig. 5A). Across animals, *shaker* rats displayed higher ataxia ratios than wild-type controls (Fig. 5B). Individual data points show non-overlapping distributions across groups, and group means differed significantly (*p* = 0.000137). Power spectral density (PSD) estimates of COM-derived speed revealed clear group differences (Fig. 5C). Wild-type rats showed smoothly decaying broadband spectra, whereas *shaker* rats exhibited increased power within the 3–8 Hz range. This difference was quantified using a normalised tremor score, which was higher in *shaker* rats than in wild-type rats (Fig. 5D; *p* = 0.000753).

**Figure 5:**
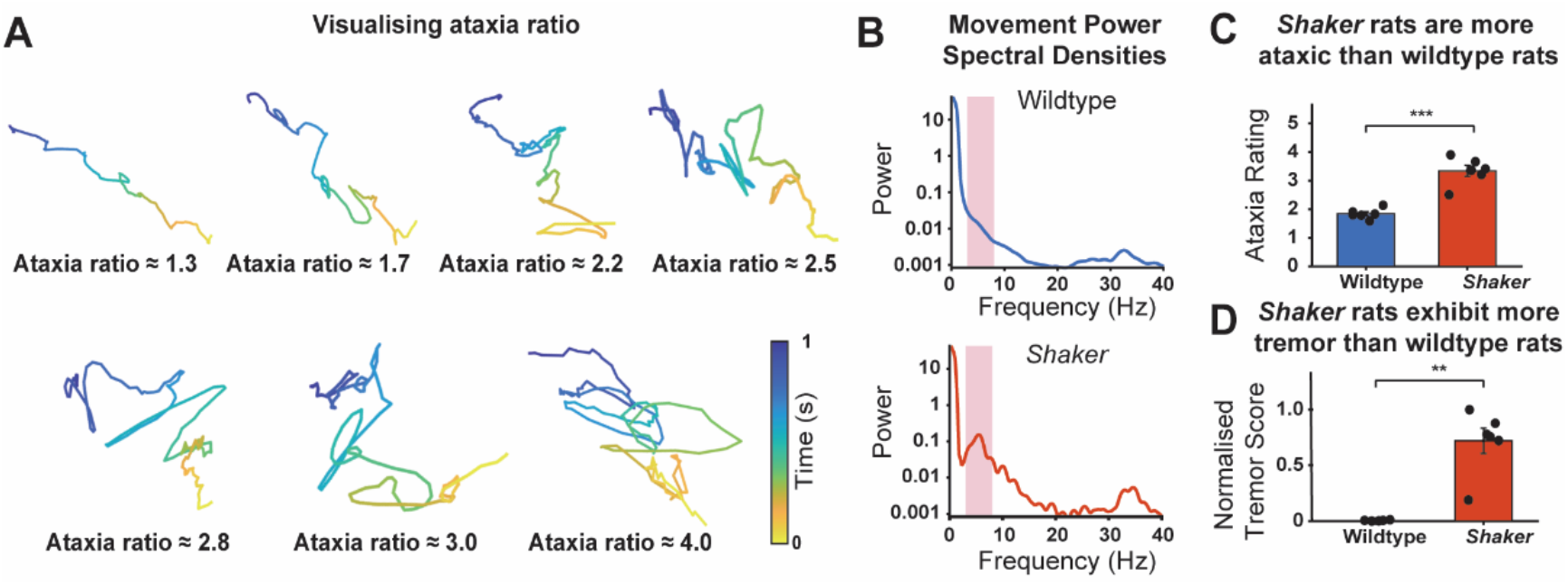
**A**. Example 1-s trajectory segments illustrating ataxia ratios ranging from approximately 1.3 to 4.0. **B**. Speed PSD estimates for representative wild-type (above) and *shaker* (below) rats. Wild-type spectra show broadband decay to 20 Hz without clear peaks, whereas *shaker* spectra display elevated power in the 3–8 Hz range (pink shaded region); **C**. Ataxia ratios for individual wild-type (n = 6) and *shaker* rats (n = 6). Data points represent per-animal means; bars show group means ± SEM. *Shaker* rats exhibited higher ataxia ratios than wild-type controls (*p* = 0.000137). **D**. Normalised tremor scores (ratio of 3–8 Hz power to total 0–20 Hz power) for wild-type and *shaker* rats. Individual points indicate per-animal values; bars show group means ± SEM. *Shaker* rats exhibit higher tremor scores than wild-type controls (*p* = 0.000753).

## 4 Discussion

In this work, we developed an open-source, low-cost force plate actometer called OpenStride, including both software and easily manufactured hardware. Currently, we have enabled OpenStride to quantify several aspects of gait and movement. Specifically, through centre-of-mass tracking in naturalistic behaviour, OpenStride quantifies speed, distance, bouts of low mobility, tremor, ataxia, and centre-*vs*-margin comparisons. The hardware described has been designed to be produced in any standard academic fabrication lab, provided the user has access to a standard large-format 3D printer and a laser cutter. At the time of publication, the total cost of manufacturing this force plate actometer is approximately $500-600 USD, assuming the user already has access to a standard computer and the aforementioned manufacturing resources. Given the open-source nature of this design, we have detailed the impact of several variables on system outcomes, and it is our intent that end users will be able to make informed decisions on how to utilise this system in a way that reflects their specific needs.

OpenStride builds off of Fowler et al.’s initial design (Fowler et al., 2001); we have previously coupled this hardware with our own custom software to quantify ataxia (Anderson et al., 2019; Anderson et al., 2025; Figueroa et al., 2023; Kuo et al., 2019; Lang et al., 2020), self-grooming behaviours (Anderson et al., 2024), and open-field assessments (Scoles 2022, bioRxiv). It was also designed to remain compatible with the above-mentioned and similar experimental paradigms, supporting quantitative analyses in studies such as pharmacological locomotor assessments (Fowler et al., 2010), investigations of dietary or developmental effects on motor activity (Levant et al., 2011; Levant et al., 2010), models of sensorimotor gating and neuropsychiatric behaviour (Godar et al., 2016), transgenic and neuroinflammatory models (Hein et al., 2012; McCarson et al., 2019), as summarised by Zarcone (Zarcone, 2022).

Several software and hardware considerations should be noted. First, while we have previously designed methods for FPA-based self-grooming quantification, we have not enabled these for OpenStride. Given that previous grooming-focussed work was carried out solely in the context of C57Bl/6 mice and dependent on machine learning-based classifiers, our enabling of this feature in OpenStride would depend on training the classifier in the context of a broader variety of models. Further, to enable this for rats, a second classifier would need to be generated and trained on several distinct rat strains for greater utility. However, given that the system saves raw data, it would be possible to design new analyses in the future and apply them to previously collected data.

Second, we note that signal-to-noise ratios will remain inferior to prior FPA designs. Original Fowler-designed FPA systems utilised higher-end Honeywell load cells at a cost of $4000 USD for the system, yielding higher-fidelity centre-of-mass recordings. Notably, given the 3D-printed nature of the OpenStride baseplate, accommodating for different sensors would be possible with moderate effort if needed. With the current OpenStride design, wild-type rat ataxia ratios averaged about 1.8, compared to ∼1.6 using an identical calculation on an original Fowler-built FPA (Anderson et al., 2019; Figueroa et al., 2023), showing the slightly higher signal-to-noise ratio of the original Fowler system.

Third, we note that temporal resolution is lower with OpenStride than with our previously reported FPA work. Previously, we interfaced load cells from Fowler FPA systems to computers using National Instruments (NI) data acquisition (DAQ) cards, with which we regularly collected centre-of-mass data at 1000 Hz. Our chosen PhidgetBridge system collects data at a maximum rate of 125 Hz, which is sufficient for most potential applications. However, modifications may be possible to enable the use of NI DAQ cards or other high-temporal-resolution input devices.

Finally, in the context of both prior FPA designs and OpenStride, signal-to-noise ratios depend on the subject’s mass, with heavier subjects resulting in reduced noise (Fig. 2). OpenStride has been designed with the intention that it can be manufactured and assembled within several days at a low cost and will function as described herein in most contexts without modification. However, given the intention of designing with low cost in mind, users intending to measure subtle changes, particularly in lightweight subjects, such as mice or very young rats, may benefit from engaging with the open-source nature of OpenStride and possibly considering use of alternative load cells.

OpenStride has been designed as an open-source, low-cost, and easily manufactured force plate actometry system. While OpenStride is currently capable of quantifying speed, distance, bouts of low mobility, tremor, ataxia, and centre-*vs*-margin comparisons, it is the intent of the authors that further capabilities will be added in the future. Given the open-source nature, numerous modifications to both hardware and software are possible, with the goal of flexibility to suit user needs. To date, force plate actometry has been used modestly in neuroscience research, with several hundred uses reported to date (Zarcone, 2022). Given a need for more naturalistic behavioural and motor quantification in numerous contexts (Cisek & Green, 2024; McCullough & Goodhill, 2021), OpenStride may serve to enable a greater uptake of force plate actometry.

## 5 Acknowledgments

The authors thank Daria Nesterovich Anderson for her generous assistance with data visualization. Further, the authors thank Stuart Kirkwood for additional advice on manufacturing and data input approaches.

## 6 Author Contributions

YY: Conceptualization, Methodology, Software, Validation, Formal Analysis, Investigation, Data Curation, Writing - Original Draft, Visualization

BC: Conceptualization, Methodology, Software, Resources, Writing - Review and Editing

MH: Conceptualization, Methodology, Software, Validation, Formal Analysis, Investigation, Writing - Review and Editing

KD: Investigation, Writing - Review & Editing, Project administration

SH: Investigation, Writing - Review & Editing, Project administration

HB: Software, Investigation, Writing - Review and Editing

KS: Investigation, Writing - Review & Editing

LH: Investigation, Writing - Review and Editing

OK: Writing - Review & Editing, Supervision

CA: Conceptualization, Methodology, Software, Validation, Forma analysis, Investigation, Resources, Data Curation, Writing - Original Draft, Writing - Review & Editing, Visualization, Supervision, Project administration, Funding acquisition

## 7 Data availability

All data associated with this manuscript will be published to GitHub along with the hardware and software files when the paper is accepted. Link will be provided here when the final version is accepted.

## 8 Additional information

The authors declare no competing interests.

